# Natural Variants of *ELF3* Affect Thermomorphogenesis by Transcriptionally Modulating *PIF4*-Dependent Auxin Response Genes

**DOI:** 10.1101/015305

**Authors:** Anja Raschke, Carla Ibañez, Kristian Karsten Ullrich, Muhammad Usman Anwer, Sebastian Becker, Annemarie Glöckner, Jana Trenner, Kathrin Denk, Bernhard Saal, Xiaodong Sun, Min Ni, Seth Jon Davis, Carolin Delker, Marcel Quint

**Author notes:** Co-first authors (Anja Raschke, Carla Ibañez).

## Abstract

Perception and transduction of temperature changes result in altered growth enabling plants to adapt to increased ambient temperature. While PHYTOCHROME-INTERACTING FACTOR4 (PIF4) has been identified as a major ambient temperature signaling hub, its upstream regulation seems complex and is poorly understood. Here, we exploited natural variation for thermo-responsive growth in *Arabidopsis thaliana* using quantitative trait locus (QTL) analysis. We identified *GIRAFFE2.1*, a major QTL explaining ~18% of the phenotypic variation for temperature-induced hypocotyl elongation in the Bay-0 x Sha recombinant inbred line population. Transgenic complementation demonstrated that allelic variation in the circadian clock regulator *EARLY FLOWERING3* (*ELF3*) is underlying this QTL. The source of variation could be allocated to a single nucleotide polymorphism in the *ELF3* coding region, resulting in differential expression of *PIF4* and its target genes, likely causing the observed natural variation in thermo-responsive growth. In combination with other recent studies, this work establishes the role of ELF3 in the ambient temperature signaling network. Natural variation of ELF3-mediated gating of *PIF4* expression during nightly growing periods seems to be affected by a coding sequence quantitative trait nucleotide that confers a selective advantage in certain environments. In addition, natural ELF3 alleles seem to differentially integrate temperature and photoperiod cues to induce architectural changes. Thus, ELF3 emerges as an essential coordinator of growth and development in response to diverse environmental cues and implicates ELF3 as an important target of adaptation.

## INTRODUCTION

In analogy to photomorphogenesis, the term *thermomorphogenesis* describes the effect of temperature on morphogenesis [1]. Hypocotyl elongation [2] and leaf hyponasty [3] belong to the most sensitive thermomorphogenic changes in plant development. Physiologically, these coordinated responses likely enhance evaporative leaf cooling [4, 5] and thus enable plants to adapt to warmth. Within the context of globally increasing ambient temperatures, it is imperative to improve our understanding of the basic processes plants employ to react to such environmental perturbations.

A major hub in the ambient temperature signaling network is the basic helix-loop-helix (bHLH) transcription factor PHYTOCHROME-INTERACTING FACTOR4 (PIF4). PIF4 protein binds to the promoters of auxin biosynthesis and response genes [6–9]. It thereby transcriptionally activates auxin responses, resulting in elongation growth. *PIF4* itself seems to be transcriptionally regulated in a temperature-dependent manner by the bZIP transcription factor ELONGATED HYPOCOTYL5 [10]. Accumulating data on PIF4 regulation from light signaling, photomorphogenesis and the circadian clock [11–13] indicate a more complex regulation of PIF4 activity on several levels.

The objective of this study was to exploit natural variation within the gene pool of *Arabidopsis thaliana* to identify additional components of the complex signaling network that plants use to adapt growth to changes in ambient temperature. Based on a quantitative genetic approach, we here show that two naturally occurring alleles of *EARLY FLOWERING3* (*ELF3*) cause a differential response in thermomorphogenesis.

## RESULTS

We previously observed extensive natural variation for the thermomorphogenic signature phenotype we termed *temperature-induced hypocotyl elongation* (TIHE; [10, 14]). To identify the underlying genetic variants, we performed quantitative trait locus (QTL) analyses based on two natural accessions from geographically distant locations. When comparing growth at 20 and 28°C, the selected accessions Bay-0 (Germany) and Sha (Tajikistan) differed significantly in several thermomorphogenic responses (Figure 1A,B; Additional file 1). Identification of the underlying genetic variants would help to improve our understanding of how ambient temperature changes are translated into growth responses.

**Figure 1.**
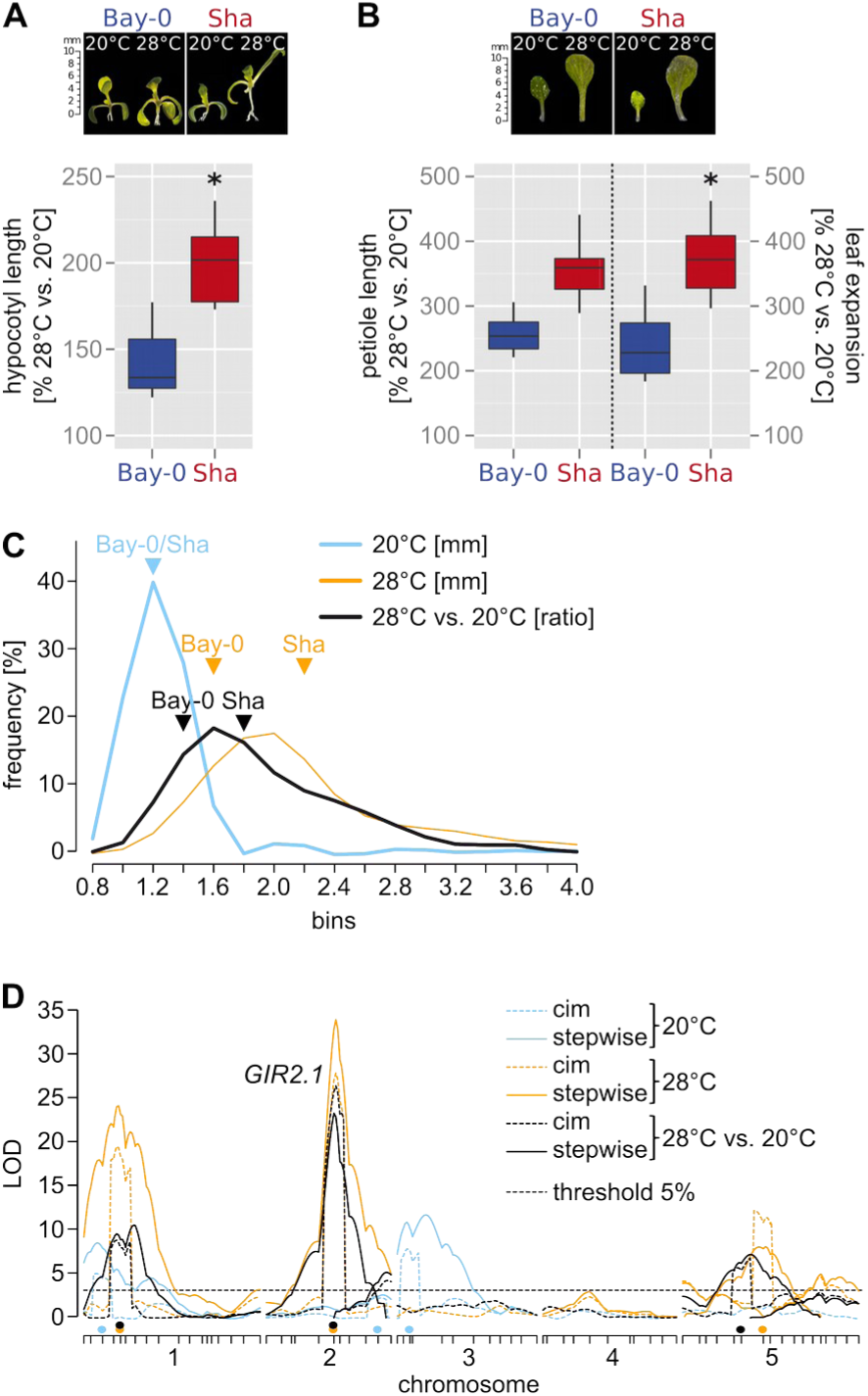
Quantitative trait locus analysis of temperature-induced growth responses in *Arabidopsis thaliana*. (A) Relative hypocotyl length (28°C/20°C in %, n=15) of 10 days-old *A. thaliana* accessions Bay-0 and Sha. (B) Relative petiole length and leaf expansion of 12 days-old seedlings. (A,B) Asterisks mark significant differences in temperature responses (P < 0.05) as assessed by two-way ANOVA of the absolute data presented in Additional file 1. (C) Frequency plot of phenotypic classes observed in a Bay-0 x Sha-derived recombinant inbred line population for hypocotyl length of 10 days-old seedlings grown at 20°C (n=400) or 28°C (n=395), and for the ratio of 28 vs. 20°C (n=387) means. Parental phenotype classes are indicated to illustrate the transgression effects within the population. (D) LOD scores (y axis) from composite interval mapping (cim) and multiple QTL mapping (stepwiseqtl) are plotted against all chromosomes (x axis). Tick marks on the x axis correspond to molecular markers in the genetic map. Colored dots on the x axis show co-variates set for composite interval mapping. Thresholds are based on 1000 permutations and an alpha of 0.05. The 28°C vs. 20°C *GIR2.1* QTL, which is the subject of this study, is highlighted.

### QTL analysis of temperature-induced hypocotyl elongation

We phenotyped a Bay-0 x Sha recombinant inbred line population [15] for the TIHE response. We grew seedlings in different ambient temperatures (10 days 20°C vs. 10 days 28°C) under a long-day diurnal cycle. QTL analysis based on composite interval and multiple QTL mapping using R/qtl was performed for hypocotyl length at either temperature alone or the ratio between hypocotyl length at 28 and 20°C (28°C/20°C). In total, we identified 14 different QTLs (Figure 1C,D, Additional files 2 and 3). Focusing on growth differences between high and low temperature (28°C/20°C ratio) identified five QTLs, which were named *GIRAFFE1/2.1/2.2/5.1/5.2* (*GIR1, GIR2.1, GIR2.2, GIR5.1, GIR5.2*), according to their respective chromosomal location. Together, the five QTLs explained ~43 % of the phenotypic variation within the mapping population. The strongest QTL, *GIR2.1* (LOD score of 23, stepwiseqtl procedure), explained ~18 % of this variation (Additional files 2 and 3), suggesting that a sizeable part of the natural variation between Bay-0 and Sha can be attributed to this locus.

To facilitate map-based cloning of *GIR2.1*, we first validated it using heterogeneous inbred families (HIFs; [16]; Figure 2A, Additional file 4). Phenotypic differences between two HIF lines, carrying either parental allele in the target region, while being otherwise genetically identical, can be attributed to genetic variation in the target QTL interval. In addition to *GIR2.1*, we also included *GIR1* and *GIR5.1* in this analysis. We were unable to validate *GIR5.1*, but observed significant differences in TIHE for the HIF lines separating the two parental alleles for *GIR1* (194-B and 194-S) and *GIR2.1* (84-B and 84-S) under long-day photoperiod (Figure 2B, Additional files 4 and 5). Due to the high impact on the phenotypic variation further analyses focused on *GIR2.1*. Here, the Sha allele conferred long hypocotyls and the Bay-0 allele conferred short hypocotyls (Figure 2B; interestingly, this situation is reversed for *GIR1* [Additional file 4]). We found that the differences in TIHE between the *GIR2.1* HIF lines did not persist under continuous light, darkness or short-day conditions (Figure 2B, Additional file 5). Hence, diurnal cycling with an extensive light phase seems to be necessary for natural variation in TIHE caused by *GIR2.1*. Furthermore, parental differences under monochromatic lights seem to be independent of *GIR2.1* (Additional file 5).

**Figure 2.**
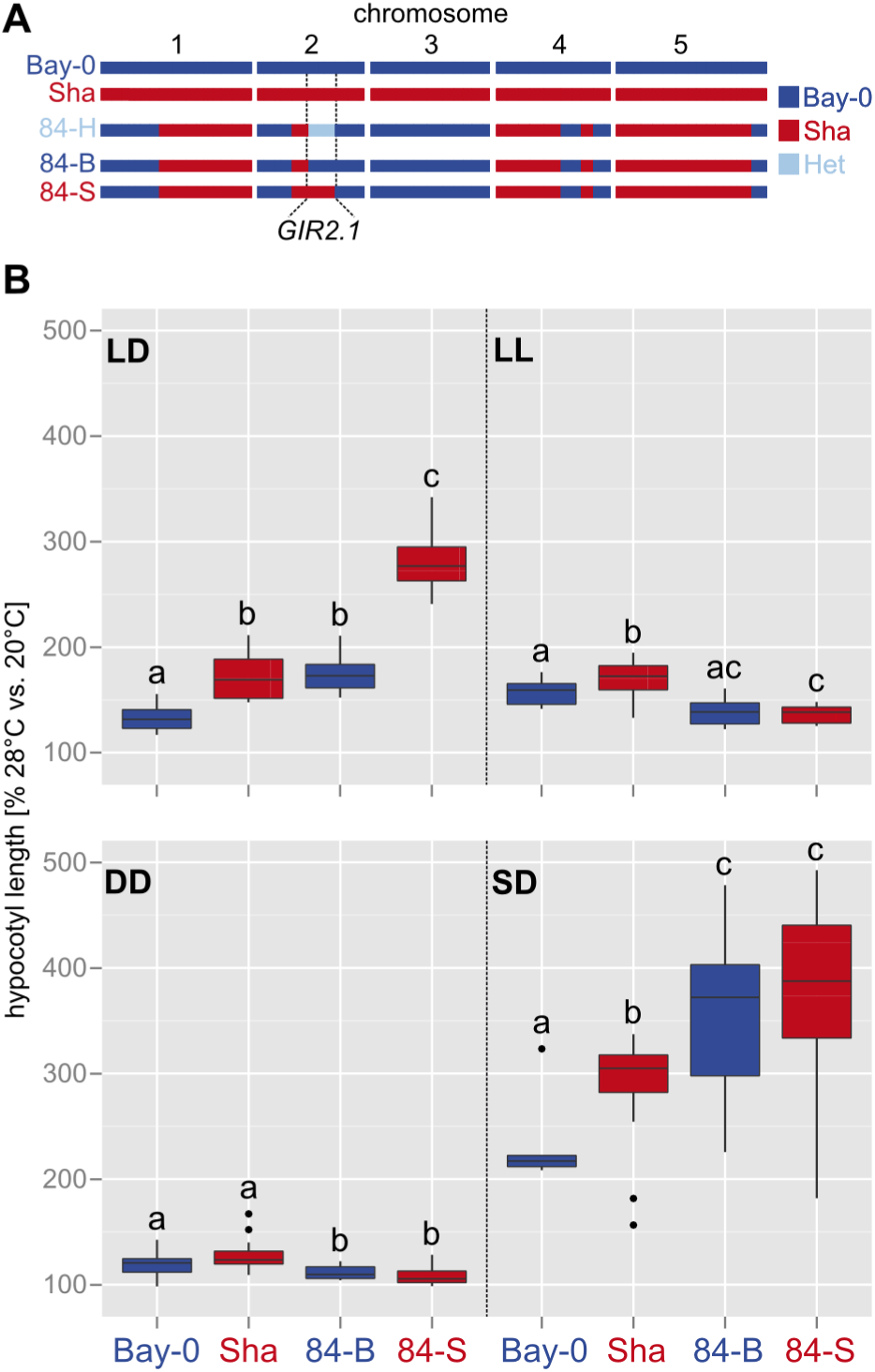
*GIR2.1* validation and photoperiod specificity. (A) Haplotype overview of the heterogeneous inbred family (HIF) 84 that segregates for Bay-0 and Sha alleles within the *GIR2.1* interval. (B) Box plots show relative hypocotyl length (28°C/20°C in %, n > 15) of seedlings grown for 8 days under long-day (LD), continuous light (LL), in darkness (DD) or short-day (SD) photoperiods. Different letters denote statistical differences in temperature responses (within one photoperiod) as assessed by two-way ANOVA (P < 0.05) of the absolute data presented in Additional file 5.

### *GIR2.1* possibly constitutes a ghost QTL

An F1 derived by crossing 84-B with 84-S showed that the long-hypocotyl phenotype inherited by the Sha allele is dominant over the Bay-0 short-hypocotyl phenotype (Additional file 5). In the process of fine-mapping the *GIR2.1* interval, genotyping of F2 and F3 recombinants with long hypocotyls (longer than 84-B) at 28°C revealed several plants for which the long-hypocotyl phenotype could be attributed to different, non-overlapping Sha intervals within the *GIR2.1* region. This indicated that the exact localization of the *GIR2.1* LOD score peak was possibly caused by two or more contributing loci. This phenomenon is frequently observed in QTL analysis and has been called *ghost* QTL [17]. Recomputing the QTL analysis with additional co-variates separated this peak into two neighbors, supporting this scenario (Additional file 6). Interestingly, Jiménez-Gómez *et al.* [18] reported a similar phenomenon for this region in the same Bay-0 x Sha population for the regulation of shade avoidance responses, which phenocopies the high temperature response. However, as we only reproducibly observed long hypocotyls for one of the two Sha intervals (chr. 2: 9,199,751-10,426,485 bp), we focused subsequent analyses on this robust interval.

### A single nucleotide polymorphism in *EARLY FLOWERING3* is underlying the *GIR2.1* QTL

Knowing that a diurnal photoperiod was a prerequisite for TIHE differences between Bay-0 and Sha (Figure 2B), we identified *EARLY FLOWERING3* (*ELF3*) as a candidate gene located in the *GIR2.1* target interval. ELF3 is a component of the circadian clock [19] that functions in the evening complex to repress growth [20] and had previously been shown to regulate hypocotyl elongation in response to shade avoidance [18]. Notably, using an elegant transgenic approach, Anwer et al. [11] recently showed that the *ELF3^Bay-0^* and *ELF3^Sha^* alleles differentially regulate period length of the circadian clock.

Consistent with a role of *ELF3* in thermomorphogenesis, we found that *elf3-4* null mutants conferred long hypocotyls in comparison to their Ws-2 wild-type (Figure 3). To investigate whether TIHE differences between the Bay-0 and Sha HIF lines can indeed be attributed to *ELF3,* we followed a transgenic complementation approach. We took advantage of the same transgenic lines generated by Anwer et al. [11] that contain either the *ELF3^Bay-0^* or the *ELF3^Sha^* allele in the *elf3-4* mutant genome (Figure 3A), which enabled us to study allele-specific TIHE effects in an independent *elf3* loss-of-function background. Figure 3B shows that transgenic lines carrying either parental allele complemented the *elf3-4* phenotype at 20°C. At 28°C, however, transgenics carrying *ELF3^Bay-0^* (*elf3-4* [*Pro_Bay-0_*:*ELF3^Bay-0^*]) repressed hypocotyl elongation significantly more than those carrying the *ELF3^Sha^* allele (*elf3-4* [*Pro_Sha_*:*ELF3^Sha^*]). This demonstrated that allelic variation in *ELF3* affects TIHE in the Bay x Sha population.

**Figure 3.**
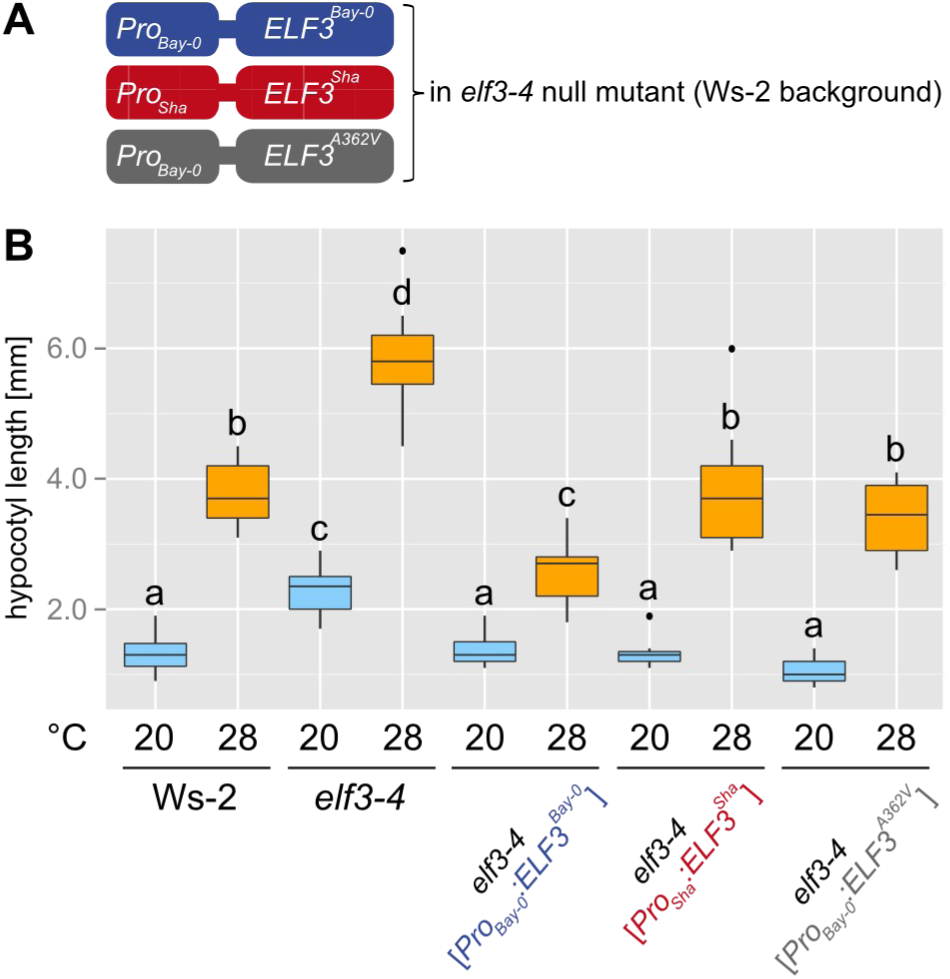
Transgenic complementation of ELF3-mediated TIHE variation. (A) Overview of transgenic constructs used for complementation of the *elf3-4* null mutation (Ws-2 background). (B) Box plot shows absolute hypocotyl length of 8 days-old seedlings grown in LD at 20 or 28°C, respectively. Different letters denote statistical differences as assessed by one-way ANOVA and Tukey HSD (P < 0.05).

Bay-0 and Sha *ELF3* variants differ (i) in a nonsynonymous SNP causing an amino acid change at position 362, encoding an alanine-to-valine transition (A362V), and (ii) in the length of a C-terminal glutamine stretch [21, 22]. Although this might depend on the genetic background [23], Tajima *et al.* [22] suggested that there was no apparent correlation between the length of the polyglutamine region and hypocotyl elongation. We therefore focused on the A362V polymorphism and investigated its potential role in conferring allelic differences in thermomorphogenesis. Again, we made use of transgenic lines in the *elf3-4* mutant background generated by Anwer et al. [11]. We inspected TIHE in transgenic *elf3-4* lines carrying either the *Pro_Bay-0_*:*ELF3^Bay-0^* allele or the *Pro_Bay-0_*:*ELF3^A362V^* allele, differing only in the A362V SNP [11]. We found that temperature-induced hypocotyls in transgenic lines carrying a valine at position 362 (as in Sha) were similar to the *ELF3^Sha^* allele and, importantly, significantly longer than those with its alanine counterpart at the same position (Figure 3B). Together, these data show that the SNP underlying the A362V change in *ELF3* causes phenotypic variation in TIHE and establishes ELF3 as a negative regulator of thermomorphogenesis.

### Differential transcriptional responses caused by natural *ELF3* variants

It has recently been shown that the evening complex of the circadian clock consisting of ELF3, ELF4, and LUX ARRHYTHMO (LUX) underlies the molecular basis for circadian gating of hypocotyl growth by directly down-regulating the expression of *PIF4* in the early evening [20]. As a result, hypocotyl elongation peaks at dawn under diurnal cycles. We, therefore, tested the hypothesis that adopts this model for ambient temperature signaling and investigated whether *PIF4* expression and possibly also the transcript levels of PIF4-regulated genes that mediate cell elongation are affected in response to elevated temperature.

To assess temperature responsiveness, we grew seedlings for 7 days at 20°C under long-day photoperiods, kept control plates at 20°C, and shifted the remaining seedlings to 28°C at lights off (t=16). Control and 28°C seedlings were subsequently harvested 4 hrs after the shift. We found that loss of *ELF3* in *elf3-4* results in up-regulation of *PIF4* transcript levels at both temperatures (Figure 4A). At 20°C, we observe that complementation of *elf3-4* with either transgenic allele (*Pro_Bay-0_*:*ELF3^Bay-0^*; *Pro_Sha_*:*ELF3^Sha^*; *Pro_Bay-0_*:*ELF3^A362V^*) restores wild-type *PIF4* levels (Figure 4A), demonstrating functionality of the constructs, but suggesting that allelic differences are absent at 20°C. This observation is consistent with similar hypocotyl length of the three lines at 20°C (Figure 3B). At 28°C, however, *PIF4* expression levels in *elf3-4* [*Pro_Sha_*:*ELF3^Sha^*] and *elf3-4* [*Pro_Bay-0_*:*ELF3^A362V^*] are significantly higher than in *elf3-4* [*Pro_Bay-0_*:*ELF3^Bay-0^*] (Figure 4A), again reflecting the hypocotyl phenotype (Figure 3B). Importantly, this expression behavior at 20 vs. 28°C explains the detection the *GIR2.1* QTL (=28°C/20°C ratio) at 28°C only, but not at 20°C (Figure 1D, Additional file 3). Together, this suggests that the natural variation observed for TIHE is attributable to temperature-dependent differences in *PIF4* expression levels caused by the A362V SNP in *ELF3^Sha^*.

**Figure 4.**
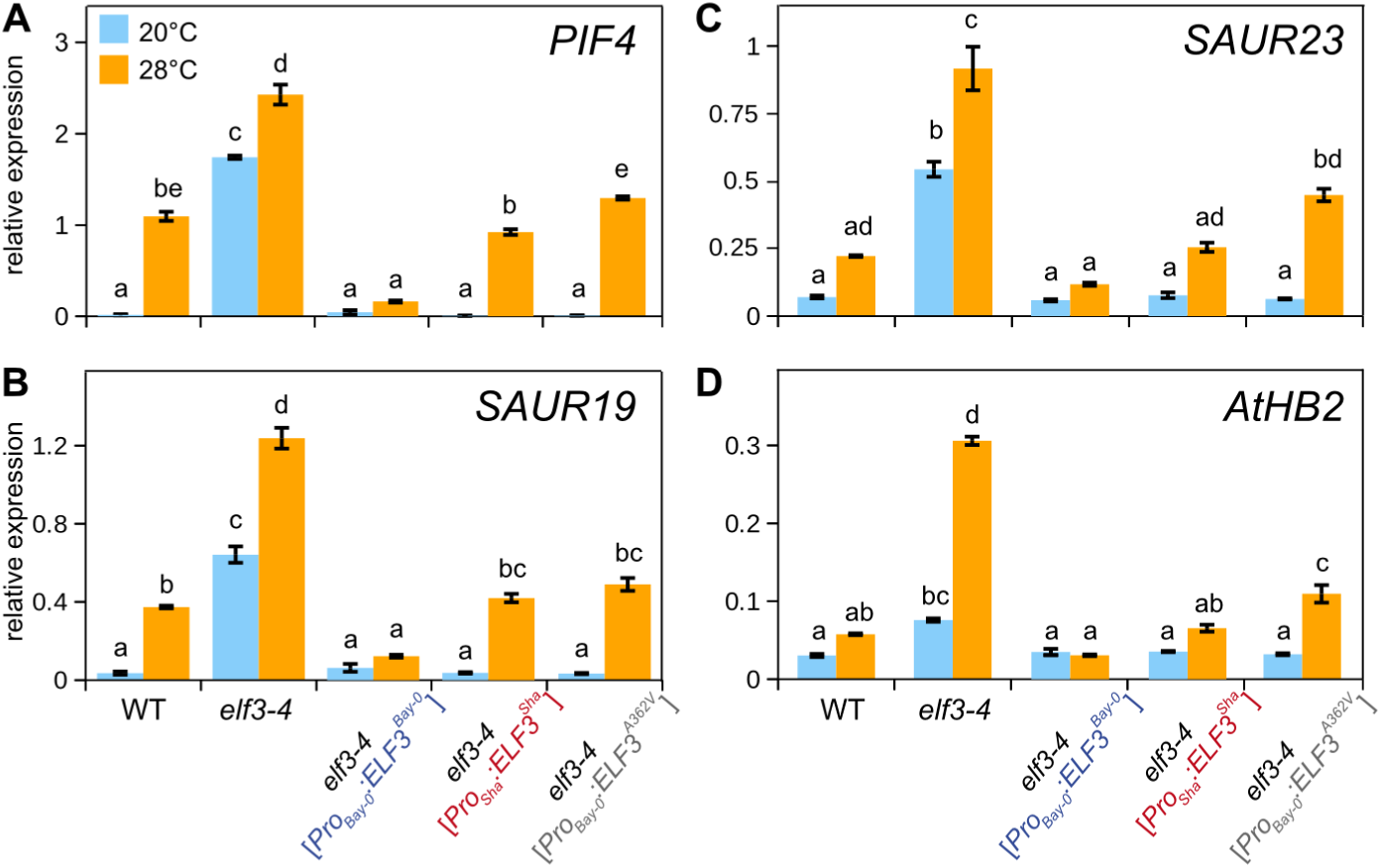
Effects of *ELF3* allelic variation on the expression of *PIF4* and auxin-responsive genes. qRT-PCR analysis of (A) *PIF4,* (B) *SAUR19,* (C) *SAUR23,* and (D) *AtHB2* expression in WT (Ws-2) and transgenic complementation lines (see Figure 3A). Seedlings were grown for 7 days at 20°C and transferred to 28°C or kept at 20°C (control). Seedlings were harvested after 4 hrs in the middle of the 8 hrs dark period. Relative expression levels of three biological replicates per treatment were assessed using *At1g13320* as control gene. Bar plots show means and SEM. Different letters denote statistical differences among samples as analyzed by one-way ANOVA and Tukey HSD test (P < 0.05).

We then connected this scenario to the level of PIF4 target genes responsible for cell elongation. Genes involved in cell elongation such as the *SMALL AUXIN UPREGULATED RNA* (*SAUR*) family or *ARABIDOPSIS THALIANA HOMEOBOX PROTEIN2* (*AtHB2*) have previously been shown to be activated by PIF4 in a temperature-dependent manner [6, 24, 25]. As Figure 4B-D shows, we found that temperature-induced expression levels of the tested genes in *elf3-4* [*Pro_Bay-0_*:*ELF3^A362V^*] were significantly higher than those in *elf3-4* [*Pro_Bay-0_*:*ELF3^Bay-0^*]. This strongly suggested that the A362V SNP causes natural variation of temperature-induced *PIF4* expression levels directly resulting in alterations of auxin-mediated cell elongation.

## DISCUSSION

Our findings shown here illustrate the power of natural variation approaches and support *ELF3* as a negative regulator of ambient temperature signaling. Physiological and gene expression data indicate that ELF3 protein might be involved in down-regulating transcript levels of the major ambient temperature signaling hub *PIF4*, and thereby affect thermoresponsive growth. Transgenic complementation assays furthermore demonstrated that a nonsynonymous SNP between the natural accessions Bay-0 and Sha significantly affects the ability of ELF3 to regulate temperature-induced *PIF4* transcript levels, its target genes, and hypocotyl elongation.

In general, different types of polymorphisms, such as nonsynonymous SNPs or expression level polymorphisms, can contribute to the expression of a particular trait [26]. In line with this phenomenon, distinct types of naturally occurring *ELF3* polymorphisms seem to contribute to hypocotyl elongation in response to different temperatures. Box *et al.* [27] recently used a different quantitative genetic approach based on the MAGIC lines [28], and elegantly showed that both protein-coding and expression level polymorphisms in *ELF3* are likely responsible for TIHE differences in natural accessions. The authors presented convincing evidence that warmth relieves the gating of growth by ELF3 at night. Specifically, ELF3 gating of transcriptional targets responds rapidly to changes in temperature by temperature-dependent binding of ELF3 to target promoters including *PIF4*. Together with Box *et al.*’s [27] non-transgenic quantitative complementation assays, our transgenic complementations unequivocally establish the role of ELF3 in thermomorphogenesis signaling.

Intriguingly, the *ELF3* QTLs in both studies were identified in different photoperiods. *ELF3* polymorphisms causal for variation within the MAGIC population were identified under short-day conditions. In contrast, our study identified the *ELF3* polymorphism under a long-day photoperiod and subsequent analysis of HIF lines showed long-day specificity (Figure 2B). Furthermore, a direct comparison of Bay-0 and Sha with two of the parental lines used in the study of Box *et al.* [27] revealed the short-day-specificity of the Sf-2 and Zu-0 alleles in promoting hypocotyl elongation (Additional file 7). This photoperiod specificity of the different natural alleles represents an interesting observation in itself requesting further investigations. Another unexpected difference between the two studies relates to the growth temperature at which the *ELF3* QTL was detected. Whereas we identified the *ELF3* QTL peak for hypocotyl growth at 28°C, but not at 20°C (Figure 1D), Box *et al.* [27] did not detect *ELF3* at high temperature but rather at 22°C standard growth conditions. This difference could be attributed to the differential integration of temperature and photoperiod by natural *ELF3* alleles. Alternatively, the genetic backgrounds and interactions with other contributing loci might be involved. In support of this, it is known that the capacity of *ELF3* to mediate growth depends on the context of the genome [23]. Hence, Box *et al.* [27] and this study confer complementary evidence for a central role of ELF3 as a major signaling hub acting upstream of PIF4 in the ambient temperature signaling network and add yet another layer to its complex regulation (Figure 5).

**Figure 5.**
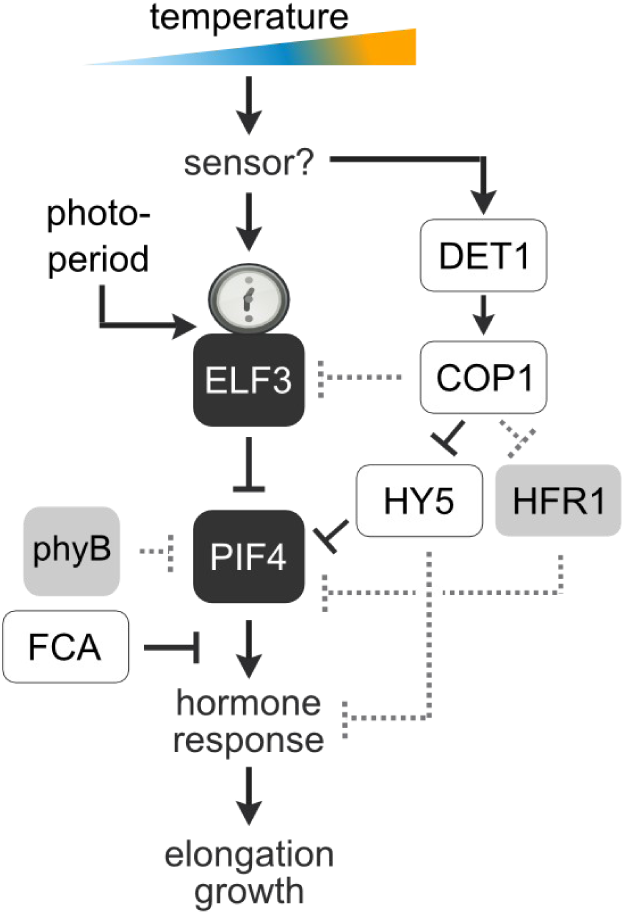
Simplified model of ambient temperature signaling. ELF3 functions as a transcriptional repressor of PIF4 and integrates temperature and photoperiod information in the regulation of thermomorphogenesis. In addition, *PIF4* regulation in response to temperature involves the regulatory components of the photomorphogenesis pathway DET1, COP1, and HY5 [10]. PIF4-mediated transcriptional regulation of target genes can be terminated by the RNA binding protein FCA, causing dissociation of PIF4 from target gene promoters [33]. Other mechanisms previously shown to contribute to PIF4 regulation in other biological contexts are depicted by gray dashed lines. These involve direct binding of DET1 to PIF4 [12], competition of PIF4 and other transcription factors for similar binding sites [34], PIF4 protein sequestration by HFR1 [35] and phyB-mediated phosphorylation and degradation of PIF4 [36].

Anwer *et al.* [11] recently identified *ELF3* as a QTL for the regulation of the circadian clock in the same Bay-0 x Sha population. In fact, they showed that the ELF3^Sha^ protein variant failed to properly localize to the nucleus and its ability to accumulate in the dark was compromised. If the same scenario holds for ambient temperature responses, then ELF3^Sha^ might fail to accumulate in the nucleus during nighttime when hypocotyl growth peaks. As a consequence of a weaker potential of ELF3^Sha^ to repress *PIF4* expression, a temperature increase could result in a much earlier activation of PIF4-mediated signaling processes during the night. Indeed, Box *et al.* [27] demonstrated that hypocotyl growth dynamics at elevated temperatures are considerably different from those at standard conditions and show a prolonged growth throughout the first night and an additional growth peak in the beginning of the dark period in subsequent nights. However, it is currently unknown in which manner temperature might affect ELF3 protein localization.

## CONCLUSIONS

In summary, remarkable progress has been made in understanding the functions of ELF3. In combination with recent studies on the role of ELF3 in the shade avoidance response [18, 21] and the circadian clock [11, 13, 29], this work contributes to understanding its role in the fine-tuned integration of a variety of environmental stimuli that in concert regulate plant growth and development (Figure 5). Natural variation in thermomorphogenesis caused by *ELF3* variants could be mediated at several levels. First, transcriptional regulation of *ELF3* itself caused by expression level polymorphisms can result in varying amounts of *PIF4*-repressing ELF3 protein [27]. In addition, coding sequence polymorphisms might affect the ability of ELF3 protein to interact with PIF4 protein and thereby inhibit its transcriptional activity, as shown by Nieto *et al.* [13]. However, it is unknown whether this protein-protein interaction is temperature-dependent and the existence of natural variation for this mechanism has yet to be reported. Lastly, nonsynonymous SNPs may affect nuclear accumulation of ELF3, which - like expression level polymorphisms described above - would result in variation of the amount of nuclear ELF3 [11] capable of transcriptionally repressing *PIF4*, and other targets. Possibly, the latter mechanism is responsible for natural variation between Bay-0 and Sha thermomorphogenesis we reported here.

In combination with the study of Box et al. [27] our work adds further insight into the essential role of ELF3 in integrating multiple signals to promote architectural changes. The photoperiod-specific function of natural ELF3 alleles could provide new avenues to elucidate the clock-mediated growth regulation in general and ELF3 mode of action specifically.

## METHODS

### Plant material

Plant material used for QTL analyses has been obtained from the Versailles Arabidopsis Stock Center: Bay-0 (accession number 41AV), Sha (236AV), heterogeneous inbred families (HIFs, 33HV84, 33HV194), Bay-0 x Sha population (33RV). Lines used for complementation assays have been described in Anwer *et al. [11].* Natural accessions Sf-2 (N6857) and (Zu-0 N6902) were obtained from the Nottingham Arabidopsis Stock Center.

### Thermo-responsive growth assays

Seeds were surface-sterilized and kept in deionized H_2_O for 3 days at 4°C before sowing. Seedlings were germinated and grown under sterile conditions and the indicated temperatures on *Arabidopsis thaliana* solution (ATS) medium [30]. Hypocotyl growth was quantified in seedlings cultivated for 8-10 days under 250 μmol m^-2^ s^-1^ white light and a long-day photoperiod (16/8) unless stated otherwise. Hypocotyl length was measured using the RootDetection software package (<u>http://www.labutils.de/</u>). Petiole length and leaf expansion were measured on 12 days-old seedlings using ImageJ. All growth assays including phenotyping of the Bay-0 x Sha population have been repeated at least three times with similar results of which one representative data set is shown.

### QTL mapping

Described QTL mapping was applied using Haley-Knott Regression [31] at 1 cM steps with the R/qtl package [32]. Logarithm of odds (LOD) score thresholds were based on 1000 permutations and an alpha error rate of 0.05. Detailed instructions on the QTL mapping procedure are found in Additional file 8. QTL mapping has been performed on all three repetitions of phenotyping of the Bay-0 x Sha mapping population independently with similar results. QTL mapping data of one representative data set are shown.

### Light response assays

Seeds were surface-sterilized, stratified at 4°C for 2 days, and dispersed on 0.8% agar (w/v)Murashige and Skoog medium. Monochromatic red, far-red, or blue light was generated with an LED SNAP-LITE (Quantum Devices, Barnereld, WI). Green light was generated from a fluorescent light bulb with a green filter. Seedling growth was measured after 4 days.

### qRT-PCR

Surface-sterilized seeds were placed on ATS medium and grown for 7 days under long-day photoperiod (16/8) and 100 μmol m^-2^ s^-1^ white light at 20°C. Temperature-induced samples were shifted to 28°C at the dusk, while control plants remained at 20°C. Samples for qRTPCR analyses were harvested in the middle of the night 4 hrs before subjective dawn. Sample preparation and qRT-PCR (including primer sequences) were performed as previously described [10].

## AUTHORS’ CONTRIBUTIONS

MQ designed the study. AR, CI, SB, AG, JT, KD, and CD performed the thermo-responsive growth assays. KKU and BS conducted the QTL analysis. AR and CI fine-mapped the GIR2.1 <u>QTL. CI</u> and CD conducted the qRT-PCR analysis. XS and MN performed the light response assays. MUA and SJD contributed materials and edited the manuscript. AR, CI, CD and MQ wrote the manuscript. All authors read and approved the manuscript.

## ACKNOWLEDGEMENTS

We thank Christine Camilleri and INRA Versailles for providing HIF lines. This work was supported by a grant from the Deutsche Forschungsgemeinschaft to MQ (Qu 141/3-1).

## ADDITIONAL FILES

**Additional file 1: Figure S1.** Temperature-induced growth responses in Bay-0 and Sha. Absolute values for (A) temperature-induced hypocotyl elongation (TIHE), (B) temperature-induced petiole elongation (TIPE), and (C) temperature-induced leaf expansion (TILE). Data correspond to the relative data presented in Figure 1.

**Additional file 2: Table S1.** Descriptive statistics of phenotypes analyzed in the Bay-0 x Sha population.

**Additional file 3: Table S2.** QTL summary statistics.

**Additional file 4: Figure S2.** Validation of the *GIR1* QTL. (A) Haplotype overview of the heterogeneous inbred family (HIF) 194 that segregates for Bay-0 and Sha within the *GIR1* interval and was used for validation and mapping of the *GIR1* QTL. (B) Box plots show relative (28°C/20°C in %) hypocotyl length of 10 days-old seedlings. Different letters denote statistical differences in temperature responses as assessed by two-way ANOVA (P < 0.05) of the absolute hypocotyl length data presented in (C). (D) Haplotype overview of the heterogenous inbred family (HIF) 214 that segregates for Bay-0 and Sha within the *GIR5.1* interval and served for the attempted validation of this QTL. (E) Box plots show relative (28°C/20°C in %) hypocotyl length of 10 days-old seedlings. Different letters denote statistical differences in temperature responses as assessed by two-way ANOVA (P < 0.05) of the absolute data presented in (F). The significant differences in TIHE observed for the parental lines Bay-0 and Sha was not reflected by the two HIF lines 214-B and 214-S that carried a Bay-0 or Sha allele within the *GIR5.1* interval, respectively. As such, the *GIR5.1* QTL could not be validated with the available genetic material.

**Additional file 5: Figure S3.** Effect of altered light conditions on *GIR2.1*-mediated hypocotyl elongation. (A) Box plots show relative (28°C/20°C in %) and absolute hypocotyl length of 10 days-old seedlings of Bay-0, Sha, and HIF lines homozygous for either Bay-0 (84-B) or Sha (84-S) in the *GIR2.1* interval. F1 plants derived from a cross of 84-B and 84-S correspond to the haplotype 84-H in Figure 2A and illustrate the dominance of the Sha over the Bay-0 allele. Different letters denote statistical differences in temperature responses as assessed by two-way ANOVA (P < 0.05) of the absolute hypocotyl length data. (B) Absolute hypocotyl length corresponding to the relative data presented in Figure 2B. (C) Hypocotyl length in monochromatic light conditions. Significant differences among Bay-0 and Sha are observed in 4 days-old seedlings grown at 20°C in constant blue (4.93 μmol m^-2^ sec^-1^), green (0.32 μmol m^-2^ sec^-1^), or red (0.89 μmol m^-2^ sec^-1^) light. These differences seem to be regulated independent of *GIR2.1* as 84-B and 84-S did not differ in their growth response. No differences among genotypes were detected in seedlings grown in far-red (0.024 μmol m^-2^ sec^-1^) light.

**Additional file 6: Figure S4.** *GIR2* constitutes a ghost QTL. Setting additional covariates in the *GIR2.1* target region separates the single peak into two linked peaks (compare with Figure 1d), indicating the potential existence of two linked loci. Tick marks on the x axis correspond to molecular markers in the genetic map of the Bay-0 and Sha mapping population. Circles on the x axis show co-variates set for composite interval mapping.

**Additional file 7: Figure S5.** Photoperiod and allele effects on ELF3-mediated TIHE. TIHE comparison of 7 days-old seedlings grown either in short day (SD) or long day (LD) photoperiods. Box plots show (A) relative (28/20°C in %) and (B) absolute hypocotyl length for Bay-0, Sha and MAGIC population parental lines Sf-2 and Zu-0 that also carry polymorphisms in *ELF3*. While Sf-2 and Zu-0 show a strong TIHE response in SD, the response for Sf-2 and Zu-0 is much weaker under LD. (A) Different letters in denote statistical differences in temperature responses as assessed by two-way ANOVA (P < 0.05) of the absolute hypocotyl length data.

**Additional file 8: Methods S1.** Estimation of heritability and QTL analysis procedure.

**Additional file 9: Supplementary Dataset 1.** Phenotypic data used for QTL mapping. This dataset has been uploaded to figshare and can be accessed via <u>http://dx.doi.org/10.6084/m9.figshare.1339892.</u>

